# Cross-Molecular Active Learning for the Discovery of Antimicrobial Polyacrylamides

**DOI:** 10.1101/2025.11.07.687243

**Authors:** Shoshana C. Williams, Gabriel Greenstein, Xinyu Liu, Alessio Fragasso, Noah Eckman, Alexander N. Prossnitz, Changxin Dong, Anna Makar-Limanov, Christine Jacobs-Wagner, Lynette Cegelski, Hector Lopez Hernandez, Eric A. Appel

**Affiliations:** Department of Chemistry, Stanford University, Stanford, CA 94305, USA; Department of Materials Science & Engineering, Stanford University, Stanford, CA 94305, USA; Department of Biology, Stanford University, Stanford, CA 94305, USA; Sarafan ChEM-H, Stanford University, Stanford, CA 94305, USA; Howard Hughes Medical Institute, Stanford University, Stanford, CA 94305, USA; Department of Chemical Engineering, Stanford University, Stanford, CA 94305, USA; Department of Microbiology and Immunology, Stanford University, School of Medicine, Stanford, CA, 94305, USA; Department of Bioengineering, Stanford University, Stanford, CA, 94305, USA; Woods Institute for the Environment, Stanford University, Stanford CA 94305, USA; Department of Pediatrics (Endocrinology), Stanford University, Stanford CA 94305, USA

## Abstract

Antimicrobial resistance poses an urgent and increasing threat to global health. The development of new antimicrobials is crucial. Synthetic copolymers are attractive as a potential solution, because they can be produced at scale and designed to mimic antimicrobial peptides and act as broad-spectrum antimicrobials capable of evading resistance mechanisms. This work leverages a cross-molecular machine learning pipeline, trained on antimicrobial peptides, to develop potent antimicrobial polymers to combat *Escherichia coli*, which were then synthesized and validated experimentally. One candidate copolymer was further characterized and shown to permeabilize the bacterial membrane, which is associated with decreased resistance. Furthermore, this copolymer demonstrated remarkable synergy in eradicating biofilm-associated *E. coli* when combined with a first-line clinical drug regimen, reducing the amount needed to eradicate bacteria in biofilms by three orders of magnitude. These results demonstrate promise for potentiating antibacterial activity of currently available antibiotics, treating serious and complicated infections, and combatting antimicrobial resistance.

## INTRODUCTION

Antimicrobial resistance (AMR) is a global emergency as antimicrobial resistant infections contribute to 4.7 million deaths annually.^1^ In the United States alone, AMR costs $4.6 billion in direct healthcare spending.^2^ The toll is even heavier in Sub-Saharan Africa and South Asia.^3^ The burden of AMR is expected to soar in the coming decades, as antimicrobial resistant strains become more prominent and existing antibiotics become increasingly ineffective. It has been estimated that by 2050, AMR will contribute to 8.2 million deaths annually.^1^

To combat this growing crisis, it is imperative to develop novel antimicrobial agents that mitigate or overcome the onset of resistance. AMR arises through several mechanisms, most commonly involving drug target modification, drug inactivation, drug uptake inhibition, and/or enhanced drug efflux.^4–6^ Host-defense peptides, or antimicrobial peptides, have gained attention in recent decades as potent, naturally-occurring antimicrobials. These peptides are typically amphiphilic, containing cationic amino acids that enhance interactions with the anionic bacterial cell surface and also containing hydrophobic amino acids to promote interaction with and disruption of the hydrophobic compartment of the bacterial cell membrane, leading to bacterial cell death.^7–12^ Notably, this mechanism simultaneously enables broad-spectrum activity and limits the development of resistance, as it does not act on a singular protein target, nor depend on intracellular access.^13,14^

Despite their promise, the clinical translation of host-defense peptides has been limited by synthetic challenges, off-target toxicity, susceptibility to proteolytic degradation, and poor pharmacokinetics.^15–17^ To address these challenges, researchers have explored the development of synthetic polymers that mimic the properties of host-defense peptides, several of which exhibit broad-spectrum activity, enhanced safety profiles, and resistance-evading properties.^18–26^ They present an affordable, effective alternative to host-defense peptides.

The further development of antimicrobial polymers is of great interest in combatting AMR and the potential chemical space available to explore is nearly infinite. This opportunity also presents a challenge as it is impossible to systematically experimentally evaluate candidates across the diverse realm of polymers to find the most potent candidates. Machine learning pipelines have demonstrated great utility in evaluating vast chemical space to identify functional molecules, including for polymer applications.^27–33^ Researchers are curating databases of antimicrobial polymers to aid in such active learning approaches; however, the utility of these databases is constrained by their relatively small size.^34,35^ In contrast, vast databases detailing the activity of host-defense peptides are available, and these have been successfully implemented to identify new, potent antimicrobial peptide candidates.^36–40^ As antimicrobial polymers are inspired by host-defense peptides and can be characterized by the same key chemical characteristics that influence peptide activity (molecular weight, charge density, hydrophobicity, etc.),^41^ we implemented an active learning pipeline leveraging peptide training data to predict antimicrobial polymers (Figure 1). We then evaluated their efficacy against *Escherichia coli* and their hemolytic activity. For one top-performing candidate, we investigated its membrane-permeabilizing activity, efficacy against biofilms, and synergy with a common, clinically used antibiotic regimen and compared these results to a previously reported antimicrobial polymer.

**Figure 1.**
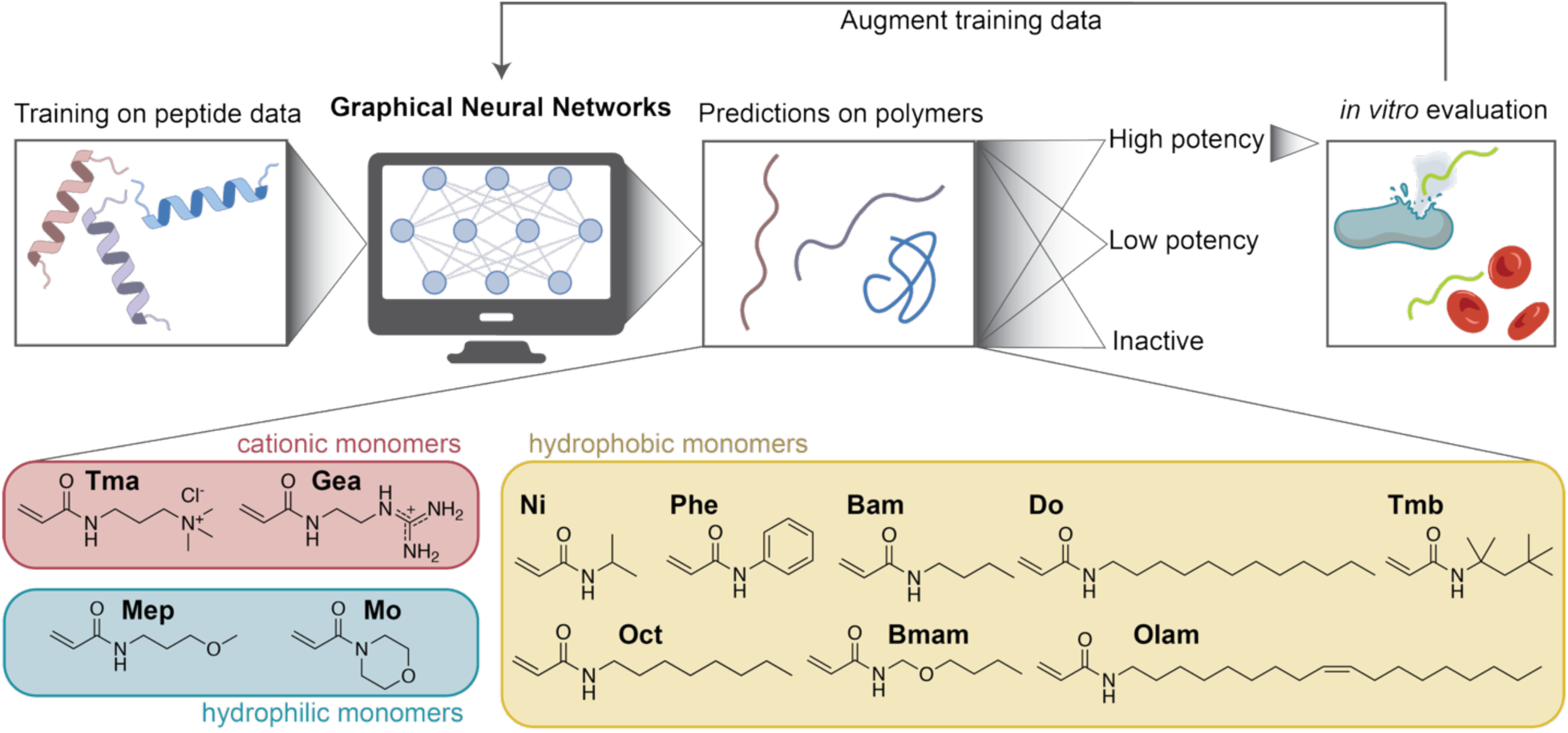
Schematic indicating the implementation of an active learning pipeline trained on peptide data to mine for potent, effective antimicrobial polymers from a large pool of synthetically accessible polyacrylamide candidates.

## RESULTS AND DISCUSSION

### Cross-molecular learning from peptides to polymers

We demonstrate that Graph Neural Networks (GNNs) trained exclusively on antimicrobial peptide data can accurately predict the antimicrobial activity of structurally distinct synthetic polymers (Figure 2; ROC-AUC: 0.91, PR-AUC: 0.92). This cross-molecular learning capability represents a significant advance over traditional approaches that require extensive training data within the target material class. By leveraging shared physicochemical properties at the monomer level, our approach enables prediction of polymer antimicrobial activity using only peptide training data, overcoming the fundamental limitation of insufficient polymer datasets in this domain.

The success of this transfer learning approach allows us to screen the vast combinatorial design space of synthetic copolymers synthesized with 2-6 monomers from a library of 12 candidate monomers: 2 hydrophilic, 8 hydrophobic, and 2 cationic. These monomers enable investigation of variations in the cationic, hydrophobic, and hydrophilic properties of the resulting copolymers, each of which are known to influence antimicrobial efficacy and safety.^42–49^ The hydrophobic monomers chosen (Figure 1) span a wide range of logP values, and these copolymer types have demonstrated broad-spectrum antimicrobial activity with low toxicity.^26^ This selection creates a combinatorial space of over 1.7 million possible synthetic polymers—far too many to screen experimentally without computational guidance.

### Model architecture and cross-molecular implementation

The development of a unified GNN model capable of predicting antimicrobial efficacy across both peptides and synthetic polymers required careful consideration of molecular representation strategies. We adopted a graph-based approach where molecules are represented as networks of interconnected monomeric units, with nodes corresponding to individual monomers and edges representing chemical bonds (peptide bonds for peptides, carbon-carbon bonds for polymers). Each node is represented by a feature vector derived from RDKit molecular descriptors, with 104 chemically relevant descriptors selected from the original 217 available descriptors based on their relevance to antimicrobial activity. This representation preserves critical structural information while enabling cross-molecular learning between chemically distinct compound classes.

A fundamental challenge in this approach stems from the inherent differences between sequence-defined peptide materials and stochastically defined synthetic polymers. While peptides have well defined molecular weights and sequences, copolymers have molecular weight distributions and compositional dispersity. To address this challenge, we implemented a stochastic sampling strategy that generates multiple possible sequences for each polymer composition (see Supplemental Information for detailed methods). Our peptide dataset comprised antimicrobial peptide data from the GRAMPA database^50^ supplemented with non-antimicrobial peptides from UNIPROT^51^ and split into train/validation/test splits. A separate holdout test set of 27 synthetic copolymers, previously synthesized and characterized in our laboratory,^26^ served as the primary evaluation benchmark for cross-molecular transfer learning performance.

We then evaluated five GNN architectures: (i) Graph Convolutional Networks (GCN), (ii) Message-Passing Neural Networks (MPNN), (iii) Graph Attention Networks (GAT), (iv) GraphSAGE, and (v) AttentiveFP. All models were implemented using the Deep Graph Library (DGL-LifeSci) framework. For each architecture, we trained 100 models using Bayesian hyperparameter optimization and selected the best performing model from each architecture for comparison (Figure S1). Based on this evaluation, we selected MPNN for the active learning pipeline, which achieved ROC-AUC: 0.91 and PR-AUC: 0.92 on the polymer holdout set (Figure 2E). The high precision-recall performance (PR-AUC: 0.92) is particularly important for experimental efficiency, as it minimizes false positives that would waste laboratory resources. The training and validation loss curves demonstrate robust convergence without overfitting, validating our model architecture and training protocol (Figure 2E, bottom panel).

### Active learning pipeline for polymer discovery

Leveraging this cross-molecular learning capability, we implemented an active learning pipeline to systematically explore the polymer design space and identify promising antimicrobial candidates. Since our initial library of antimicrobial polymers was small and our model’s performance was validated on only a subset of the chemical space, we expected model predictions to provide noisy labels. Building on this assumption, we used the trained GNN model to power an active learning loop, whereby model predictions guided the selection of screening candidates.

Our active learning strategy comprised three distinct sampling phases that progress systematically from exploration to exploitation (Figure 2B-D): (i) diversity sampling to ensure broad coverage of the chemical space, (ii) uncertainty sampling to train our models with samples in which it had the least confidence, and (iii) exploitation sampling to identify high-efficacy candidates. This systematic progression from exploration to exploitation ensures comprehensive coverage of the design space while efficiently identifying high-value targets.

We conducted three rounds of diversity sampling, including two initial rounds using terpolymers (6 polymers total, 4 of which were previously published^26^) and one expanded round incorporating 4-monomer copolymers (3 additional polymers). We employed a MaxMin diversity [RDKit MaxMin Sampler] sampling algorithm that maximizes the minimum pairwise distance between selected polymers in a chemical feature space (Figure 2B).

We conducted the two rounds of uncertainty sampling (10 polymers each) using an ensemble-based approach that identifies regions where multiple trained models disagree in their predictions (Figure 2C). We trained multiple independent neural network models and measured the disagreement between their predictions using KL-divergence from the ensemble mean. Polymers with high inter-model disagreement were prioritized for experimental validation, as these represent regions where the models are most uncertain and additional data would be most valuable for improving predictive performance. This strategy successfully identified polymers in chemically ambiguous regions of the design space, enabling targeted refinement of model decision boundaries (see Supplemental Information for detailed methods).

**Figure 2.**
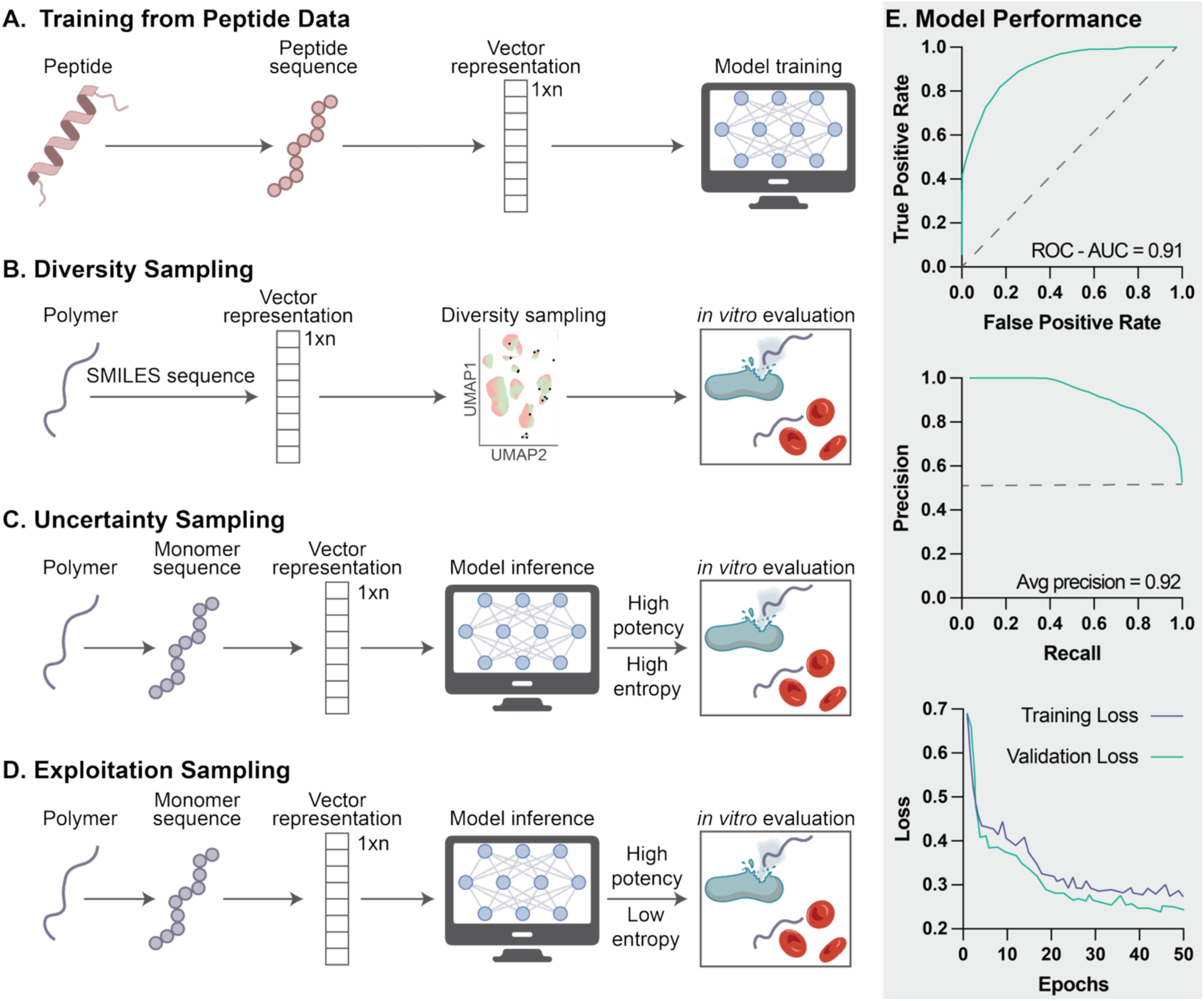
Schematic indicating the active learning workflow, including **A.** model training, **B.** diversity sampling, **C.** uncertainty sampling, and **D.** exploitation sampling. **E.** Model performance after training, evaluated on a test set of known antimicrobial polymers.

We then conducted one round of exploitation sampling (5 polymers) designed to identify high-efficacy candidates by classifying polymers with a minimum inhibitory concentration (MIC) less than 100 μg/mL (Figure 2D). We selected candidates with the highest model confidence with lowest ensemble disagreement (low KL-divergence).

The complete active learning pipeline synthesized and characterized 34 polymers across six sampling rounds. Our final polymer library included an additional 23 rationally designed and previously screened polymers, for a total of 57 polymers.

### Efficacy and safety of candidate copolymers

The copolymers identified through diversity sampling, uncertainty sampling, and exploitation sampling were synthesized using RAFT polymerization from commercially available or readily synthesized monomers (Figure 1 lists monomers and their abbreviations). Each candidate copolymer was prepared by statistical copolymerization of mixtures of 2-6 monomers and named according to those monomers and their weight percent in the composition (Table 1). Each copolymer was analyzed by NMR and GPC (Figure S4; Figure S5).

**Table 1.** Composition of synthesized polyacrylamides.

| **Polymer** | **Target DP<sup>a</sup>** | **Target M<sub>w</sub> (kDa)<sup>a</sup>** | **% Conversion<sup>b</sup>** |
| --- | --- | --- | --- |
| Gea<sub>40</sub>Olam<sub>10</sub>Mo<sub>50</sub> | 70 | 11.8 | 92.5 |
| Gea<sub>80</sub>Phe<sub>5</sub>Mo<sub>15</sub> | 70 | 12.6 | 84.3 |
| Gea<sub>5</sub>Phe<sub>25</sub>Mo<sub>50</sub>Mep<sub>20</sub> | 70 | 10.1 | 95.9 |
| Tma<sub>50</sub>Do<sub>5</sub>Mo<sub>40</sub>Mep<sub>5</sub> | 70 | 12.0 | 99.3 |
| Tma<sub>70</sub>Phe<sub>20</sub>Olam<sub>5</sub>Mep<sub>5</sub> | 70 | 13.0 | 78.2 |
| Tma<sub>75</sub>Do<sub>25</sub> | 70 | 15.0 | 98.5 |
| Gea<sub>10</sub>Olam<sub>30</sub>Mo<sub>60</sub> | 70 | 12.3 | 98.9 |
| Tma<sub>55</sub>Ni<sub>25</sub>Phe<sub>20</sub> | 70 | 11.2 | 99.5 |
| Tma<sub>55</sub>Bam<sub>40</sub>Oct<sub>5</sub> | 70 | 11.5 | 97.8 |
| Tma<sub>17</sub>Phe<sub>23</sub>Oct<sub>5</sub>Mo<sub>55</sub> | 70 | 10.7 | 99.9 |
| Tma<sub>45</sub>Gea<sub>10</sub>Phe<sub>20</sub>Olam<sub>25</sub> | 70 | 14.5 | 73.2 |
| Tma<sub>35</sub>Gea<sub>20</sub>Ni<sub>35</sub>Tmb<sub>10</sub> | 70 | 11.0 | 96.8 |
| Tma<sub>35</sub>Phe<sub>35</sub>Olam<sub>10</sub>Mo<sub>20</sub> | 70 | 12.1 | 92.6 |
| Tma<sub>38</sub>Gea<sub>17</sub>Tmb<sub>8</sub>Mo<sub>37</sub> | 70 | 12.1 | 99.2 |
| Tma<sub>30</sub>Gea<sub>20</sub>Ni<sub>20</sub>Olam<sub>25</sub>Mo<sub>5</sub> | 70 | 13.0 | 94.1 |
| Tma<sub>90</sub>Do<sub>10</sub> | 70 | 14.7 | 56.8 |
| Tma<sub>80</sub>Phe<sub>15</sub>Do<sub>5</sub> | 70 | 13.7 | 89.7 |
| Tma<sub>85</sub>Gea<sub>5</sub>Phe<sub>5</sub>Olam<sub>5</sub> | 70 | 14.4 | 84.3 |
| Tma<sub>70</sub>Ni<sub>20</sub>Do<sub>5</sub>Mep<sub>5</sub> | 70 | 12.3 | 93.5 |
| Tma<sub>60</sub>Phe<sub>20</sub>Olam<sub>5</sub>Mo<sub>15</sub> | 70 | 12.8 | 98.6 |
| Tma<sub>40</sub>Phe<sub>5</sub>Do<sub>5</sub>Mo<sub>50</sub> | 70 | 11.6 | 98.9 |
| Tma<sub>70</sub>Gea<sub>10</sub>Oct<sub>15</sub>Tmb<sub>5</sub> | 70 | 14.0 | 77.9 |
| Tma<sub>65</sub>Do<sub>5</sub>Bam<sub>5</sub>Mo<sub>20</sub>Mep<sub>5</sub> | 70 | 12.7 | 98.9 |
| Tma<sub>25</sub>Gea<sub>5</sub>Phe<sub>15</sub>Bam<sub>30</sub>Mo<sub>25</sub> | 70 | 10.6 | 94.3 |
| Tma<sub>70</sub>Phe<sub>5</sub>Olam<sub>10</sub>Bmam<sub>10</sub>Mep<sub>5</sub> | 70 | 13.9 | 81.9 |
| Tma<sub>35</sub>Gea<sub>5</sub>Ni<sub>10</sub>Phe<sub>5</sub>Do<sub>30</sub>Mep<sub>15</sub> | 70 | 14.7 | 72.9 |
| Tma<sub>30</sub>Do<sub>15</sub>Bam<sub>15</sub>Oct<sub>10</sub>Mo<sub>30</sub> | 70 | 12.8 | 98.8 |
| Tma<sub>15</sub>Gea<sub>10</sub>Ni<sub>5</sub>Do<sub>25</sub>Tmb<sub>10</sub>Mep<sub>35</sub> | 70 | 14.0 | 73.5 |
| Tma<sub>25</sub>Ni<sub>10</sub>Do<sub>5</sub>Mo<sub>60</sub> | 70 | 12.7 | 99.2 |
| Tma<sub>20</sub>Gea<sub>15</sub>Ni<sub>10</sub>Do<sub>10</sub>Olam<sub>10</sub>Mep<sub>35</sub> | 70 | 10.6 | 70 |

<sup>a</sup>Target DP and Target M<sub>w</sub> are theoretical values.
<sup>b</sup>Percent conversion was determined by <sup>1</sup>H NMR analysis.

These copolymer materials were then evaluated for their potential antimicrobial efficacy through determination of the MIC against *E. coli* grown in MHB at a cell density of 5 x 10^5^ CFUs/mL (Figure 3; Table S1). In total, five copolymers were identified with experimentally confirmed MIC values below 100 μg/mL (range: 32-64 μg/mL). This result demonstrates the utility of ensemble-based approaches for both exploration and exploitation in active learning pipelines (see Supplemental Information for detailed methods).

To evaluate the potential for undesirable mammalian toxicity, hemolysis was measured for each copolymer (Figure 3; Table S1). Nineteen polymers demonstrated very low hemolysis, with HC_50_ > 4000 μg/mL. Notably, two polymers (Tma_75_Do_25_ and Tma_70_Phe_5_Olam_10_Bmam_10_Mep_5_) demonstrated both impressive efficacy as described above, with MICs of 64 µg/mL, while also exhibiting HC_50_ values among the highest reported for antimicrobial polymers.^42,49,52–54^ Interestingly, these copolymers were identified through uncertainty sampling rather than exploitation sampling, which may indicate dispersity in the strength of each model, as greater agreement among them did not necessarily correspond to greater activity.

**Figure 3.**
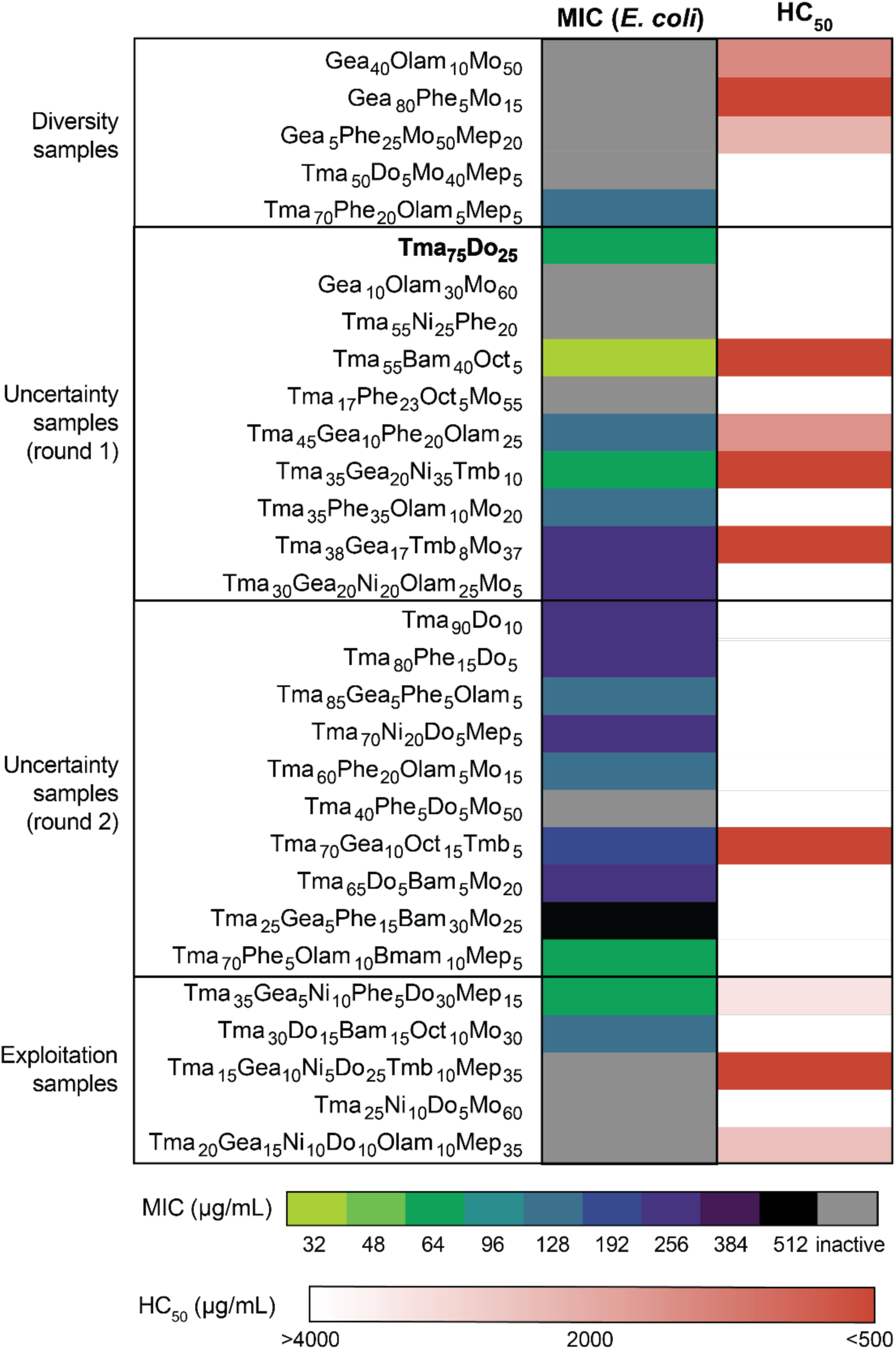
Safety and efficacy of novel polyacrylamide derivatives. (Left) Heat map indicating the MIC of each copolymer against *E. coli*. (Right) Heat map indicating their hemolytic activity.

### Copolymer activity against the bacterial membrane

One copolymer candidate (Tma_75_Do_25_) was selected for further evaluation. This candidate was compared to a previously reported copolymer identified through rational design (Tma_59_Do_31_Mep_10_; previously labeled L-Do_31_Mep_10_).^26^ *E. coli* were incubated with the DNA-intercalating probe SYTOX Green in the presence of each antimicrobial copolymer. A rapid and dramatic increase in SYTOX Green fluorescence was observed upon the addition of the antimicrobial copolymer (Figure 4A), indicating the copolymers induce damage to the inner membrane of the bacteria. The damaged inner membrane, which is impermeable to SYTOX Green when intact, enables the SYTOX Green’s intracellular access and subsequent intercalation into DNA. This result is general across both copolymers, and no statistically significant difference between the polymers was observed in the SYTOX Green permeability (p = 0.14).

**Figure 4.**
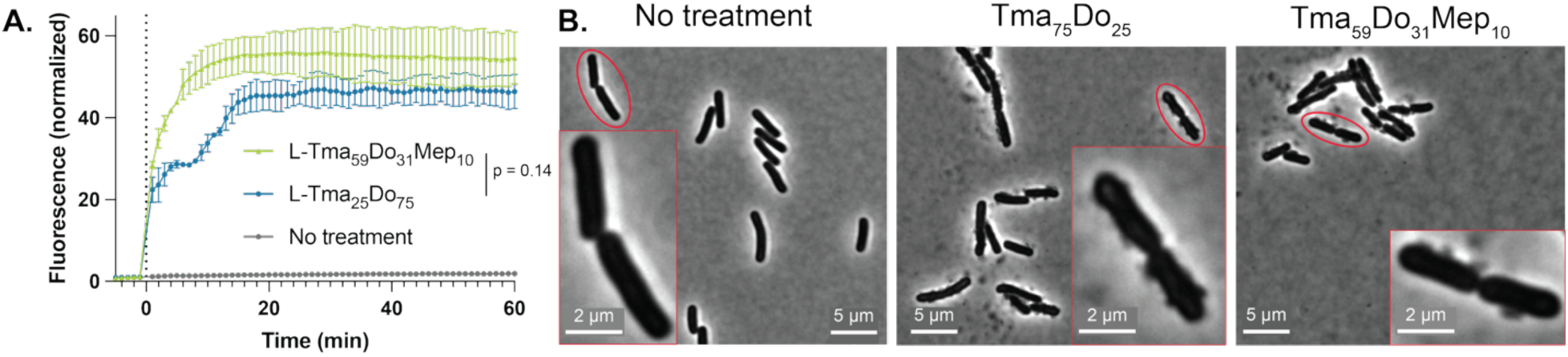
Mechanism of antimicrobial copolymers. **A.** Membrane permeabilization assay for *E. coli* treated with each polymer at 2x MIC, using the fluorescent probe SYTOX Green, monitored continuously by plate reader. **B.** Optical microscopy of *E. coli* after treatment with each polymer at 2x MIC.

Additionally, copolymer-treated *E. coli* were examined by phase-contrast light microscopy (Figure 4B). Ruffling was observed on the surface of the bacterial cells after treatment with each polymer relative to untreated *E. coli*. Together, these results demonstrate that these antimicrobial copolymers disrupt the surface of the bacteria and compromise the cell membranes.

### Antimicrobial copolymers eradicate biofilm-associated bacteria and potentiate clinically-used antibiotics

Biofilm infections are notoriously difficult to eradicate with many clinically available antibiotics. This is in part because bacteria in a biofilm are protected by extracellular polymeric matrix, and they can be less metabolically active and therefore less susceptible to antibiotics that target cell growth.^55^ As such, biofilms can cause persistent or chronic infections in humans, with limited treatment options.^56^ New antimicrobials capable of combating biofilms are therefore highly desirable.

To test the efficacy of our leading copolymers against biofilms, we determined their minimum biofilm eradication concentration (MBEC) against *E. coli* grown in M63 medium with glucose. For each copolymer, the MBEC was 512 µg/mL (Table 2, Figure 5, Figure S7), which falls within the expected therapeutic window and far below the HC_50_ (above 4000 µg/mL for each). By contrast, an antibiotic regimen commonly used as a first-line treatment for urinary tract infections (trimethoprim-sulfamethoxazole, abbreviated as TMP-SMX)^57^ demonstrates an MBEC of 256 µg/mL, which may approach the limit of its therapeutic window, as its LD50 for injection in mice is 700 mg/kg (Table 2, Figure 5, Figure S7).^58^ This result demonstrates the utility of these antimicrobial copolymers against biofilm infections.

**Table 2.** Efficacy of copolymers and TMP-SMX against biofilm-associated bacteria, alone and in combination.

| Treatment | TMP-SMX | Tma<sub>75</sub>Do<sub>25</sub> | Tma<sub>59</sub>Do<sub>31</sub>Mep<sub>10</sub> |
| --- | --- | --- | --- |
| MBEC alone, $\mu\text{g/mL}$ ($\mu\text{M}$) | 256 (984.9) | 512 (21.9) | 512 (42.2) |
| MBEC for TMP-SMX-Tma<sub>75</sub>Do<sub>25</sub> combo, $\mu\text{g/mL}$ ($\mu\text{M}$) | 0.125 (0.5) | 64 (2.7) | n/a |
| MBEC for TMP-SMX-Tma<sub>59</sub>Do<sub>31</sub>Mep<sub>10</sub> combo, $\mu\text{g/mL}$ ($\mu\text{M}$) | 0.25 (1.0) | n/a | 64 (5.3) |

**Figure 5.**
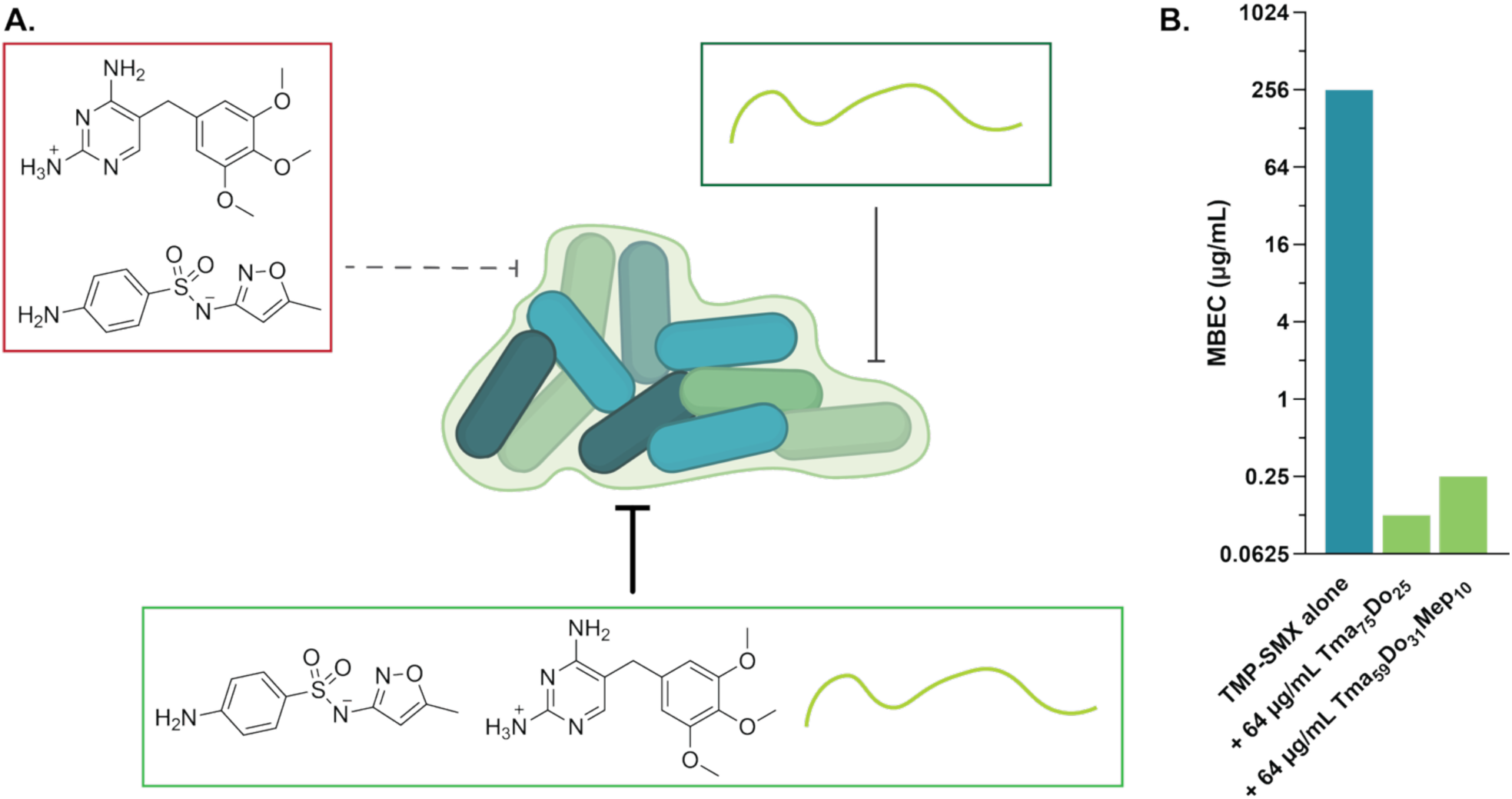
Anti-biofilm efficacy of copolymers. **A.** Schematic indicating the anti-biofilm and antibiotic-potentiating activity of copolymers. **B.** Comparison of MBEC for TMP-SMX antibiotic treatment alone and in combination with each polymer.

The copolymers were then explored as adjuvants to investigate their potential to synergize with TMP-SMX. Each exhibited remarkable synergy in combination with TMP-SMX (Table 2, Figure 5, Figure S7). For the combination of Tma_75_Do_25_ and TMP-SMX, the polymer MBEC was 64 µg/mL (corresponding to 1/8^th^ of its MBEC value alone) and the TMP-SMX MBEC was 0.125 µg/mL (corresponding to 1/2048^th^ of its MBEC value alone). For the combination of

Tma_59_Do_31_Mep_10_ and TMP-SMX, the polymer MBEC was 64 µg/mL (corresponding to 1/8^th^ of its MBEC value alone) and the TMP-SMX MBEC was 0.25 µg/mL (corresponding to 1/1024^th^ of its MBEC value alone). The fractional inhibitory concentration indices (FICIs) are 0.1255 or 0.1260 for the combinations with Tma_75_Do_25_ and Tma_59_Do_31_Mep_10_ respectively, indicating exceptionally strong synergy. This remarkable finding demonstrates the ability of the copolymers to dramatically improve the efficacy of a first-line antibiotic treatment, bringing it back to its therapeutically relevant dosing. The copolymers show potential to expand the scope of existing antibiotics in clinical applications.

## CONCLUSION

New antimicrobial agents are urgently needed to address the growing crisis of antimicrobial resistance. Towards that goal, this work introduces a cross-molecular active learning pipeline to identify polyacrylamide-based copolymers with antimicrobial activity against *E. coli*. We implemented GNNs trained on antimicrobial peptides to predict the properties of copolymer compositions, demonstrating the potential for cross-molecular training. We screened predicted copolymers to identify compositions that are effective and non-toxic, and selected one top candidate, Tma_75_Do_25_, for further investigation. This copolymer was compared to a previously reported polymer, Tma_59_Do_31_Mep_10_. Fluorescent nucleic acid staining and microscopy imaging demonstrated that these copolymers damage the bacterial cell surface and permeabilize the inner membrane. Furthermore, we revealed tremendous synergy and potentiation a first-line antibiotic regimen, TMP-SMX, in combination with each copolymer. Remarkably, each combination tested reduced the TMP-SMX concentration needed to eradicate bacteria in biofilms by three orders of magnitude, positioning it in the highly therapeutically attractive sub-micromolar range.

Future directions will expand on this promising strategy and should focus on incorporating multi-objective optimization into the machine learning pipeline to allow for simultaneous optimization on multiple goals, such as potency against multiple strains of bacteria and favorable safety profiles. In addition, while the copolymer design space described here is combinatorially large at 1.7 million compositions, it is constrained to 12 monomers representing a narrow scope of possible copolymer chemistries. There is enormous potential to expand the chemistry and activities of the materials screened. The incorporation of diverse backbone architectures, alternative polymerization mechanisms, and broader monomer libraries will be essential for developing the most powerful, truly generalizable predictive models. There is also potentially a need to implement sophisticated methods for modeling molecular weight dispersity^59^ and compositional dispersity^60^ to enable more rigorous modeling of *in silico* polymerizations in a machine learning pipeline. Additional work should further explore the efficacy of the reported copolymer candidates against more bacterial species, both *in vitro* and *in vivo*. This work has demonstrated that these copolymers are effective antimicrobials with low hemolysis and the potential to rescue existing antibiotics to combat dangerous biofilm-associated infections. Further efforts toward their translation could significantly broaden our arsenal of effective agents to combat against antimicrobial resistance.

## AUTHOR INFORMATION

### Author contributions

The study was conceptualized by S.C.W., H.L.H., and E.A.A with contributions from L.C. and C.J-W. The active learning pipeline was developed by H.L.H, G.G., and N.E. Experiments were conducted by S.C.W., A.F., X.L., C.D., A.M.L., and A.N.P. The original draft was written by S.C.W.

## ACKNOWLEDGEMENTS

We thank Dr. Emily Meany and Dr. Jerry Yan for kindly providing murine blood for hemolysis assays. We thank Dr. Carolyn Jons for her advice regarding statistical characterization.

