## Supplemental Methods and Data for "Cross-Molecular Active Learning for the Discovery of Antimicrobial Polyacrylamides"

*Materials*

All solvents were used as received: *N,N*-dimethylformamide (DMF; Sigma-Aldrich), hexane (Sigma-Aldrich), acetone (Aldon Corporation), diethyl ether (Fisher Scientific), ethyl acetate (EtOAc; Sigma-Aldrich), dichloromethane (DCM; Sigma-Aldrich) methanol (MeOH; Sigma-Aldrich), ($D_2O$; Sigma-Aldrich), chloroform-d ($CDCl_3$; Sigma-Aldrich), and dimethyl sulfoxide-d6 (DMSO-d6; Sigma-Aldrich).

Acryloyl chloride (Sigma-Aldrich), triethylamine (TEA; Sigma-Aldrich), *N*-octylamine (TCI Chemicals), oleylamine (Sigma-Aldrich), 1,3-Bis(tert-butoxycarbonyl)-2-methyl-2-thiopseudourea (Sigma-Aldrich), ethylenediamine (Sigma-Aldrich), and trifluoroacetic acid (TFA; Sigma-Aldrich or TCI Chemicals) were used as received.

Monomers *N*-phenylacrylamide (Phe; Sigma-Aldrich), *N*-dodecylacrylamide (Do; TCI Chemicals), *N*-butylacrylamide (Bam; TCI Chemicals), *N*-(Butoxymethyl)acrylamide (Bmam; TCI Chemicals), *N*-(1,1,3,3-Tetramethylbutyl)acrylamide (Tmb; TCI Chemicals) were used as received. *N*-isopropylacrylamide (Ni) was recrystallized hexanes. 4-acryloylmorpholine (Mo; Sigma-Aldrich) and *N*-(3-methoxypropyl)acrylamide (Mep; Sigma-Aldrich) were filtered through basic alumina before use. (3-Acrylamidopropyl)trimethylammonium chloride (Tma; 75 wt. % in $H_2O$, Sigma-Aldrich) was washed thrice with an equal volume of EtOAc and placed under vacuum for two minutes to remove residual EtOAc.

Azobisisobutyronitrile (AIBN; Sigma-Aldrich) was recrystallized from MeOH and dried under vacuum before. 4-((((2-Carboxyethyl)thio)carbonothioyl)thio)-4-cyanopentanoic acid (CTA; Sigma-Aldrich) was used as received.

Bacterial strains were purchased from Microbiologics. Mueller-Hinton broth (MHB; BD Difco), tryptic soy media (Sigma-Aldrich), and agar (Sigma-Aldrich) were autoclaved before use.

Fluorescent images were acquired using a Nikon Ti2-E inverted microscope equipped with a Photometrics Prime BSI back-illuminated sCMOS camera (2048x2048 pixels, 6.5 μm pixel size), a 100X Plan Apo 1.45NA Ph3 oil objective (with type N immersion oil by Nikon), a Lumencore Spectra III LED (Light Emitting Diode) engine, a Perfect Focus System (PFS), temperature-controlled Okolab enclosure, and a polychroic mirror (FF-409/493/596-Di02 for by Shemrock) combined with a triple-pass emitter (FF-1-432/523/702-25 by Shemrock).

*Dataset preparation and molecular representation*

<u>Antimicrobial peptide dataset:</u> Our training dataset comprised antimicrobial peptide sequences from the GRAMPA database, which aggregates 51,345 peptides from multiple databases (APD, DADP, DBAASP, DRAMP, and YADAMP).[1] We retained only peptides reporting minimum inhibitory concentrations (MICs) against *Escherichia coli*, excluded sequences with unusual modifications, and collapsed duplicate entries by taking the geometric mean of their MIC values. This yielded 5,154 unique positive AMPs with lengths ranging from 2 to 190 amino acids (mean 22.6 ± 12.6 aa) and MIC values spanning 0.023-3,200 μM (mean 60.2 ± 213.3 μM). Non-antimicrobial peptides were obtained from UniProt,[2] selecting reviewed (Swiss-Prot) entries

tagged as generic peptides with lengths between 1-40 amino acids, excluding any annotated with "Antimicrobial" keywords. After removing duplicates and non-canonical residues, we obtained 5,917 negative samples (mean length 24.9 ± 10.8 aa), assigned placeholder MIC values of 9,999 μM. The final peptide corpus contained 11,003 distinct sequences for model training and validation. The dataset was partitioned into training (70%), validation (15%), and test (15%) splits using stratified sampling to maintain class balance across splits.

Synthetic polymer dataset: A holdout test set of 27 synthetic copolymers, previously synthesized and characterized in our laboratory, served as the primary evaluation benchmark for cross-molecular transfer learning performance. These polymers were synthesized from a library of 12 monomers representing diverse chemical functionalities: 2 hydrophilic, 8 hydrophobic, and 2 cationic monomers. The polymers span compositions from 2-6 monomers with experimentally determined antimicrobial activities against *E. coli*.

Graph representation: Both peptides and polymers were represented as molecular graphs where nodes correspond to monomeric units (amino acids for peptides, chemical monomers for polymers) and edges represent covalent bonds. Node features were derived from RDKit molecular descriptors computed using rdkit.Chem.Descriptors, generating 217 descriptors per SMILES string. From these, we selected 104 descriptors deemed chemically relevant to antimicrobial activity by excluding descriptors that primarily scale with chain length (SIZE), are redundant with retained descriptors (REDUN), represent simple atom counts with little variance (COUNT_dup), encode max/min versions of properties already captured by richer descriptors (EXTREMA), or count functional groups rarely present in peptides and antimicrobial polymers (FRAG_rare). Edge features encoded bond types and connectivity patterns, with 79 edge features retained after removing zero-variance descriptors across the combined peptide and polymer datasets.

Polymer sequence generation: To address the inherent polydispersity of synthetic copolymers, we implemented a stochastic sampling strategy that generates multiple possible sequences for each polymer composition. Each polymer is defined by its monomeric composition (monomer identities and molar ratios) and average degree of polymerization (DP). To be compatible with the sequence-based format of peptide data, we sampled sequences from each polymer distribution, constraining the sample means to match the corresponding experimental composition and DP. This approach captures the intrinsic variability of linear polymer synthesis. During training, we sampled 50 sequences per polymer composition. For evaluation, we generated 200 sequences per polymer and determined final predictions through majority voting across all realizations.

### *Graph neural network architecture and training*

Model architecture: We evaluated five GNN architectures: Graph Convolutional Networks (GCN), Message-Passing Neural Networks (MPNN), Graph Attention Networks (GAT), GraphSAGE, and AttentiveFP. GCN applies convolution directly over node features, treating each neighbor equally, while MPNN updates nodes by aggregating messages from neighboring nodes and their connecting edge features. GAT introduces self-attention to learn which neighbors deserve more weight, and AttentiveFP combines local message propagation with a global attention mechanism to capture both short- and long-range interactions. Notably, GCN and GAT only use node features, whereas MPNN and AttentiveFP also incorporate edge information. All models were implemented using the Deep Graph Library (DGL-LifeSci) framework with consistent hyperparameter optimization protocols.

Hyperparameter optimization: For each architecture, we trained 100 models using Bayesian hyperparameter optimization with Tree-structured Parzen Estimator (TPE) sampling. Optimization targeted validation set performance over 50 iterations, with early stopping based on validation loss plateauing.

Model selection: The best-performing model from each architecture was selected based on validation set ROC-AUC (Figure S1). MPNN was chosen for the active learning pipeline based on superior cross-molecular transfer performance (ROC-AUC: 0.91, PR-AUC: 0.92 on the polymer holdout set).

*Active learning pipeline*

Design space definition: The polymer design space encompassed copolymer combinations synthesized from 12 candidate monomers (structures shown in Figure 1 of main text). Each copolymer in the design space contained 2-6 monomers at 5 wt% compositional intervals. Some rational design was employed in further defining the design space: each polymer was at least 10 wt% cationic and 10 wt% hydrophobic. For the uncertainty and exploitation sampling, the total concentration of Gea was limited to minimize unintended crosslinking, and Gea and Bmam were not copolymerized, due to the chemical incompatibility of Bmam with the Gea deprotection reaction. In total, over 1.7 million unique compositions were utilized.

Diversity sampling: We employed MaxMin diversity sampling to ensure broad coverage of the chemical space. The algorithm maximizes the minimum pairwise distance between selected polymers in a molecular descriptor space derived from RDKit molecular fingerprints. Distance calculations used Tanimoto similarity coefficients, with selection prioritizing compositions that maximize the minimum distance to previously selected samples.

Query-by-Committee uncertainty sampling: We implemented a Query-by-Committee (QBC) approach using ensemble disagreement to quantify prediction uncertainty. Three independent MPNN models were trained with different random initializations and hyperparameter configurations to create a diverse ensemble.

For each polymer composition, model disagreement was quantified using Kullback-Leibler (KL) divergence between individual model predictions and the ensemble mean. The ensemble mean positive and negative probabilities were calculated across all models, with a tolerance factor ($1\times10^{-6}$) applied to prevent numerical instabilities from log(0) calculations.

The KL divergence for each model's prediction p relative to the ensemble mean was computed as:

$$KL(p||q) = p\cdot\log_2(p/q\_pos) + (1-p)\cdot\log_2((1-p)/q\_neg)$$

where q_pos and q_neg represent the ensemble mean positive and negative probabilities, respectively. The final uncertainty score was the mean KL divergence across all ensemble models.

To focus on genuinely uncertain predictions while avoiding statistical outliers, we applied quartile-based filtering. Polymers were ranked by KL divergence scores, and only those from the fourth quartile (Q4) were considered for selection. Outliers were removed using the 1.5×IQR method, excluding samples beyond Q3 + 1.5×IQR. From this filtered set of high-uncertainty candidates, diversity sampling was applied using molecular descriptors to ensure broad chemical space coverage within the uncertain region.

Exploitation sampling: For exploitation sampling, we employed the same ensemble approach but focused on extracting the probability of polymers belonging to the lowest MIC bin (<100 μg/mL) from multiclass predictions. We selected polymers with high predicted probability for the lowest MIC class (>0.8) and low KL divergence scores (<0.1), indicating strong ensemble consensus for high-efficacy predictions.

*Synthesis of N-octylacrylamide (Oct)*

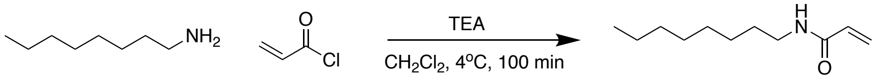

*N*-octylacrylamide was prepared by an adaptation of previously described methods.[3,4] Briefly, *N*-octylamine (992 uL, 6 mmol) and TEA (920 uL, 6.6 mmol) were dissolved in anhydrous DCM (60 mL) and cooled in an ice bath. The mixture was placed under a nitrogen atmosphere and stirred as acryloyl chloride (536 uL, 6.6 mmol) was added dropwise over 1 h. The mixture was stirred for an additional 40 minutes. It was then washed with saturated ammonium chloride (100 mL) followed by saturated sodium bicarbonate (100 mL) and brine (100 mL). The product was dried over anhydrous magnesium sulfate, and the solvent was removed by rotary evaporator. The product (Oct) was isolated as a waxy white solid with 97% yield. NMR spectra were recorded on a Bruker Avance Neo 500 MHz NMR spectrometer by collecting 32 scans with a 2 second relaxation delay.

*Synthesis of oleylacrylamide (Olam)*

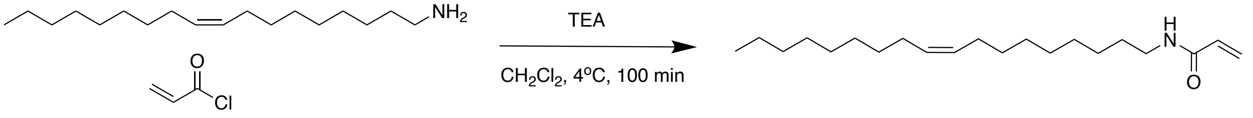

Oleylacrylamide was prepared by previously reported methods.[4] Briefly, oleylamine (1.974 mL, 6 mmol) and TEA (920 µL, 6.6 mmol) were dissolved in anhydrous DCM (60 mL) and cooled in an ice bath. The mixture was placed under a nitrogen atmosphere and stirred as acryloyl chloride (536 uL, 6.6 mmol) was added dropwise over 1 h. The mixture was stirred for an additional 40 minutes. It was then washed with saturated ammonium chloride (100 mL) followed by saturated sodium bicarbonate (100 mL) and brine (100 mL). The product was dried over anhydrous magnesium sulfate, and the solvent was removed by rotary evaporator. The product (Olam) was isolated as a pale yellow oil with 87% yield. NMR spectra were recorded on a Bruker Avance Neo 500 MHz spectrometer by collecting 32 scans with a 2 second relaxation delay.

*Synthesis of di-Boc-guanidinium ethyl acrylamide (Gea)*

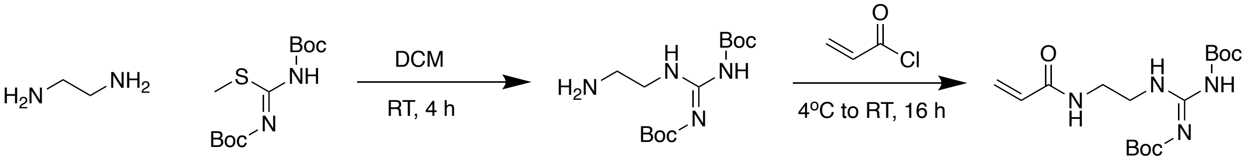

Di-Boc-guanidinium ethyl acrylamide was prepared by an adaptation of previously reported methods.[5] Briefly, 1,3-Bis(*tert*-butoxycarbonyl)-2-methyl-2-thiopseudourea (7.260 g, 25 mmol, 1 eq.) in DCM (50 mL) was added dropwise over 45 mins to a solution of ethylenediamine (5.0 mL, 75 mmol, 3 eq.) in DCM (50 mL) in an oven-dried 250 mL round-bottom flask, stirring at 500 rpm. Following complete addition, the reaction was left to stir at room temperature for 4 h. The solution was washed with $H_2O$ (2 x 50 mL) and brine (1 x 50 mL), the organic layer was dried over $Na_2SO_4$ and the DCM was removed under vacuum to yield crude 2-[1,3-Bis(*tert*-butoxycarbonyl)guanidine]ethylamine as a white solid (7.07 g, 93% yield).

2-[1,3-Bis(*tert*-butoxycarbonyl)guanidine]ethylamine (1.5 g, 4.96 mmol, 1 eq.) was transferred to an oven-dried 100 mL RBF equipped with a stir bar. TEA (761 µL, 5.46 mmol, 1.1 eq.) was added, and the solution was cooled in an ice bath for 15 minutes while stirring. The reaction mixture was equipped with an $N_2$ balloon. Acryloyl chloride (403 µL, 4.96 mmol, 1 eq.) was diluted in 1 mL DCM, and this solution was added dropwise by syringe. The balloon was removed. The reaction was covered with aluminum foil and allowed to stir and warm slowly to RT overnight with stirring at 400 rpm. The mixture was washed with saturated NaHCO3. The aqueous layer was extracted with 50 mL DCM. The organic layers were combined and dried over $Na_2SO_4$, and the solvent was removed under vacuum, yielding di-Boc-guanidinium ethyl acrylamide as an off-white solid (1.29 g, 73% yield).

*Polymerization conditions*

Statistical copolymerizations were carried out by RAFT polymerization. For each polymer, [CTA]/[monomers] = 70. [AIBN]/[CTA] was between 0.2 and 0.5 for each entry. Generally, [AIBN]/[CTA] = 0.5 for entries containing Morph and 0.2 for other copolymers. The monomers (total mass of 0.5 g), CTA, and initiator were dissolved in a mixture of DMF and water to a final volume of 4 mL in a 20 mL scintillation vial equipped with a PTFE/silicone septum. The mixture was sparged with nitrogen for 10 minutes and heated at 65 °C overnight with stirring. The mixture was cooled to room temperature and exposed to air.

Polymers containing Gea were deprotected to remove the Boc groups by mixing the crude polymerization mixture with TFA (750 µL). The mixture was equipped with a vent needle and allowed to stir at 300 rpm at room temperature for 1-2 h.

Polymers were precipitated two to three times in acetone or a combination of ether and hexane. The precipitated polymers were flash-frozen and dried by lyophilization.

*Polymer characterization*

NMR spectra were recorded on a Bruker Avance Neo 500 MHz spectrometer by collecting 128 scans with a 5 second relaxation delay. $M_n$, $M_w$, and dispersity values were determined via GPC implementing PEG standards after passing through an SEC column (Resolve Mixed Bed Low divinylbenzene(DVB) (Jordi Labs)) in a mobile phase of DMF with 1 wt% $LiBF_4$ at 50 °C and a flow rate of 1.0 mL/min (Dionex UltiMate 3000 pump, degasser, and auto-sampler (Thermo Fisher Scientific)).

To computationally probe the compositional dispersity of top candidate polymers, a previously described program, Compositional Drift, was utilized.[6] Briefly, this program predicted composition for a sample of 1500 simulated polymer chains from a pool of 200,000 monomers using the Mayo-Lewis model of monomer addition. Reactivity ratios of each monomer were assumed to equal 1. The predicted set of polymers was analyzed to determine the coefficient of variance in the molar ratio of each monomer.

*Minimum Inhibitory Concentration (MIC) assay*

The MIC of each polymer against *E. coli* ATCC 25922 was determined by broth microdilution method, adapted from the Clinical and Laboratory Standards Institute (CLSI) guidelines.[7] Briefly, *E. coli* ATCC 25922 was cultured in Mueller-Hinton broth (MHB) overnight at 37 °C with shaking at 200 rpm. The absorbance of the culture was read at 600 nm, and the culture was diluted to 1 x $10^6$ colony-forming units (CFUs)/mL. A two-fold dilution series of 50 µL of polymer solution in media was prepared in a 96-well, U-bottom polypropylene plate. The treatment media was prepared by spiking MHB with a stock solution of each polymer at 20.48 mg/mL in water to reach the desired final concentration. To each well were added 50 µL of the diluted bacterial culture, so each well contained a final concentration of 5 x $10^5$ CFUs/mL. All samples were run in duplicate. Each plate also contained sterile controls (without bacteria) and growth controls (without polymer). The plates were incubated statically at 37 °C for 18-24 h. After incubation, each well was mixed by pipetting to resuspend any bacterial growth, and 75 μL from each well were transferred to a clear, flat-bottom 96-well plate. The absorbance was measured at 600 nm on a BioTek Synergy H1 microplate reader. The MIC was defined as the lowest concentration before bacterial growth increased by at least 0.1 absorbance units.

*Hemolysis assay*

Red blood cells (RBCs) were collected by centrifugation of whole blood at 500 xg for 5 min. The plasma was removed, and the RBCs were washed twice with 150 mM sodium chloride and once with PBS. They were pelleted again and resuspended in PBS. The sample was diluted 1:50 in PBS. To each well of a 96-well plate was added 180 μL of diluted RBCs and 20 μL of antimicrobial agent stock solution at 10x the desired concentration. Samples were run in duplicate. Triton X-100 at a final concentration of 1% was used as a positive control, and water was used as a negative control. The plate was incubated statically at 37°C for 1 h. Intact RBCs were pelleted by centrifugation at 1,000 xg for 5 min using a Thermo Scientific Sorvall Legend XTR centrifuge. 100 μL of the supernatant were transferred from each well to a clear, flat-bottom 96-well plate, and the absorbance was measured at 540 nm. The percentage of hemolysis was calculated as follows:

Hemolysis (%) = [($Abs_{450\ nm}$ of the treated sample – $Abs_{450\ nm}$ of the negative control)/($Abs_{450\ nm}$ of positive control – $Abs_{450\ nm}$ of negative control)] × 100%.

The $HC_{50}$ was determined through interpolation after fitting the percentage of hemolysis to a five-parameter asymmetric sigmoidal function using GraphPad Prism software.

*Fluorescence-based membrane integrity assay*

SYTOX Green was used to evaluate the integrity of the cell membranes of *E. coli* in the presence of antimicrobial agents. Briefly, a stationary phase culture of *E. coli* ATCC 25922 was diluted 1:25 in MHB and grown at 37 °C with shaking at 200 rpm to until the absorbance at 600 nm ≈ 0.6. The bacteria were subsequently pelleted by centrifugation at 10,000 rpm for 5 min at room temperature using a Thermo Scientific Sorvall Legend Micro 21R centrifuge. They were washed once with HEPES/glucose buffer (5 mM HEPES and 5 mM glucose). They were pelleted again and resuspended to a final optical density of 1 in HEPES/glucose buffer. The cells were aliquoted into a black-walled 96-well plate, and SYTOX Green was added to a final concentration of 1 μM (from a 20x stock solution prepared in HEPES/glucose buffer). The fluorescence was measured in a microplate for 5 min (excitation 490 nm, emission 520 nm). The plate was removed, and the antimicrobial agents were added at the desired concentrations (from 20x stock solutions prepared in HEPES/glucose buffer). The total volume in each well was 150 μL. The fluorescence was measured for 1 h at room temperature. Statistical analysis comparing the fluorescence curve for each polymer was conducted with GraphPad Prism software using a 2-way ANOVA.

*Light microscopy*

Phase-contrast light microscopy was performed as previously reported,[9] using the *E. coli* strain ATCC 25922. Briefly, the cells were grown to stationary phase in MHB medium at 37 °C. To obtain exponentially growing cells, the culture was then diluted by at least 10,000-fold in fresh MHB and grown at 37 °C until it reached an optical density at 600 nm ($OD_{600}$) of 0.2-0.4.

Polymer solutions were prepared by diluting a 20 mg/mL stock solution in fresh MHB and mixed 1:1 with the cell solution by pipetting, yielding a final polymer concentration of 2xMIC and $OD_{600}$ = 0.1. As a control, the same step was performed with fresh MHB instead of the polymer solution. Samples were incubated for 30 min at 37 °C in a thermomixer (500 rpm), and 0.5 µL was spotted onto a 1% (w/v) agarose pad. This pad was prepared by adding agarose powder in fresh MHB. The suspension was then gently warmed to dissolve the agarose. Exposure time for phase-contrast images was 200 ms with 100% diascopic light intensity, and images were acquired at room temperature.

*Minimum Biofilm Eradication Concentration (MBEC) assay*

*E. coli* ATCC 25922 biofilms were formed by overnight static incubation at 37 °C of 100 µL of inoculum (1 x $10^7$ CFU/mL in M63 minimal media with 0.8% w/v glucose) in the inner wells of a sterile PVC plate. To help with growth consistency of the biofilms and evaporation, 150 µL of PBS was added to the outer row and column of each plate. After 24 h of incubation, the growth media was then carefully removed from the inner wells by pipette, and the wells were rinsed three times with 120 µL of sterile PBS to remove planktonic cells. After rinsing, 150 µL treatment media containing 2-fold serial dilutions of the antimicrobial compound in M63 + 0.8% glucose was added to the plate. The treatment media was prepared by spiking M63 + 0.8% glucose with a stock solution of the antimicrobial agent: $Tma_{75}Do_{25}$ were prepared at 10.24 mg/mL in water, $Tma_{59}Do_{31}Mep_{10}$ at 10.24 mg/mL in 50:50 water/DMSO, and TMP-SMX at 30 mg/mL in DMSO. The plate was incubated statically for an additional 24 h at 37 °C. The treatment media was then removed and the wells were again rinsed three times with 160 µL PBS. Fresh recovery media (150 µL M63 with 0.8% glucose) was added and the plate was incubated statically for an additional 24 h at 37 °C. Then, 15 µL of 0.02% resazurin solution was added to each well and incubate for 30 min. The MBEC was read as the lowest treatment concentration where no bacterial growth occurred, as determined visually according to the color change of the resazurin dye.

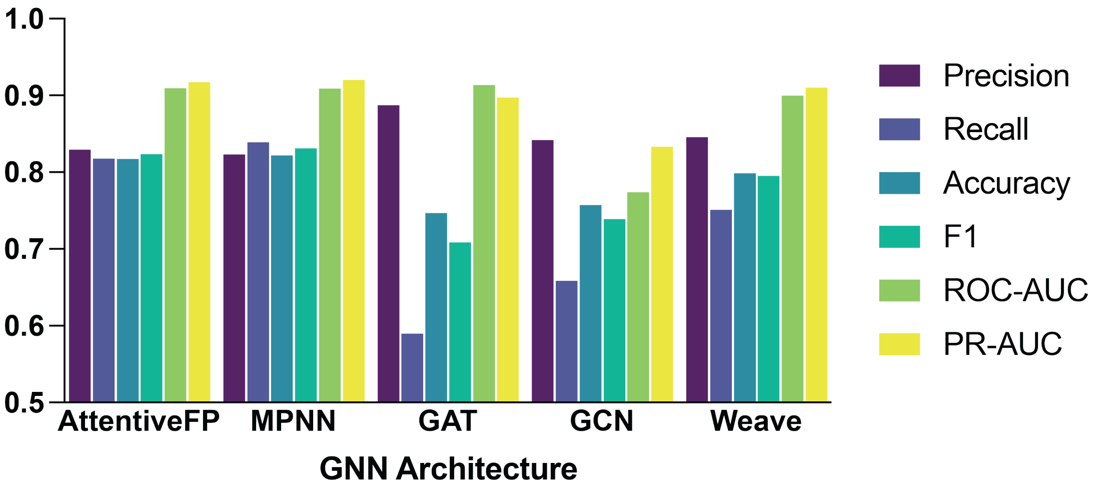

**Figure S1. Comparison of top-performing GNN architectures.** The best-performing model from each architecture on a polymer test set with precision > 0.8. Performance defined as the average of precision, recall, F1, accuracy, ROC-AUC (area under the receiver operating characteristic curve), and PR-AUC (area under the precision-recall curve).

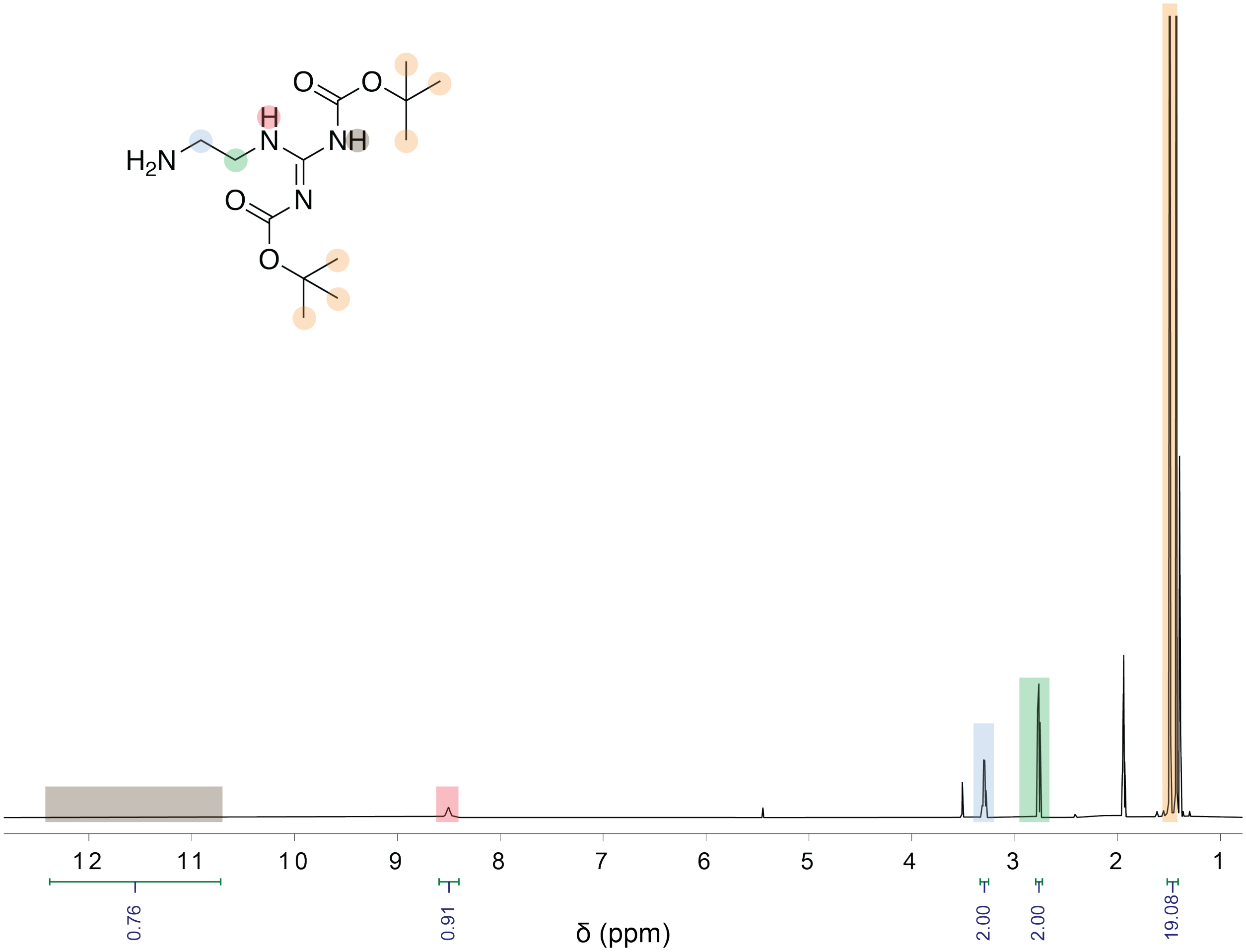

**Figure S2. $^1$H NMR Spectrum of 2-[1,3-Bis(*tert*-butoxycarbonyl)guanidine]ethylamine.** Measured in deuterated acetonitrile.

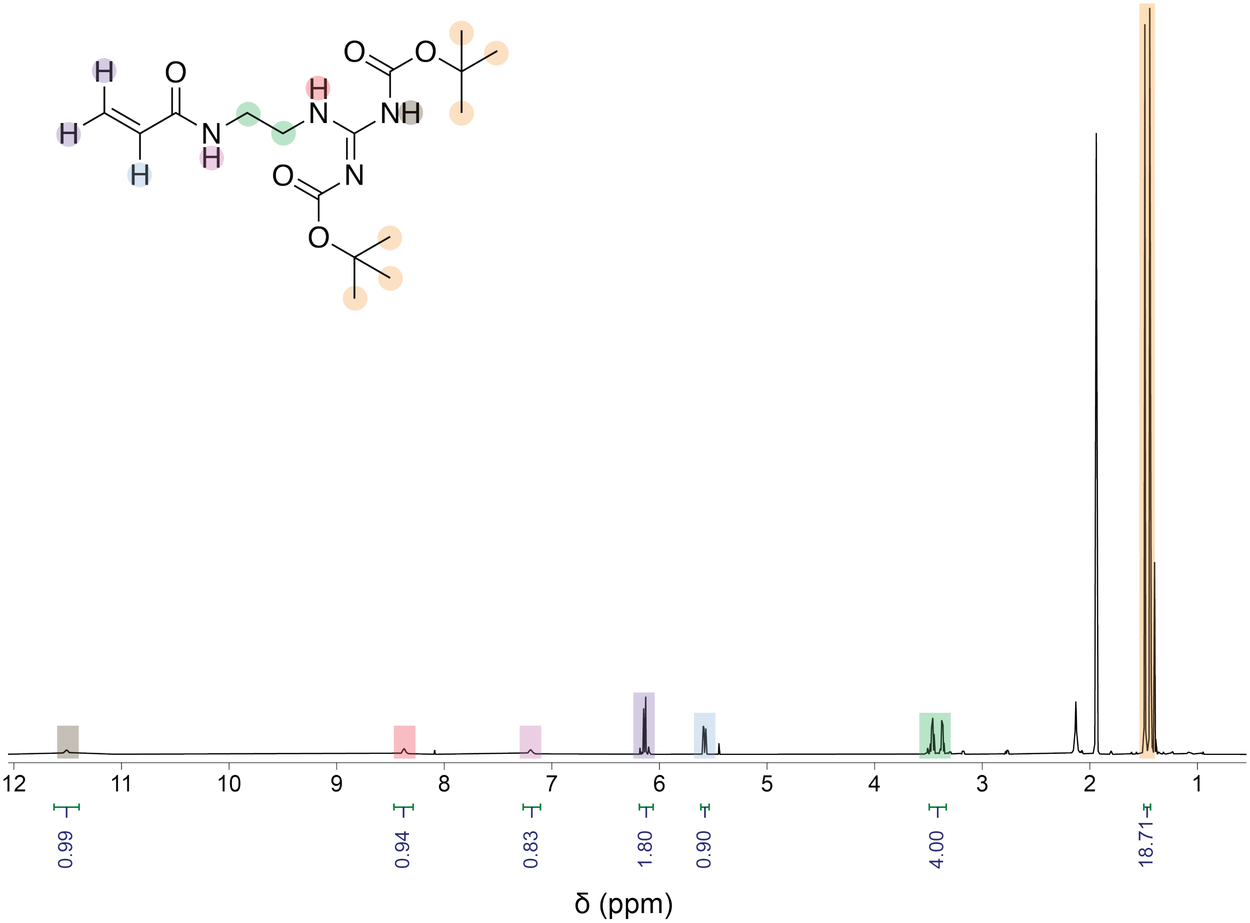

**Figure S3. [1]H NMR Spectrum of di-Boc-guanidinium ethyl acrylamide (Gea).** Measured in deuterated acetonitrile.

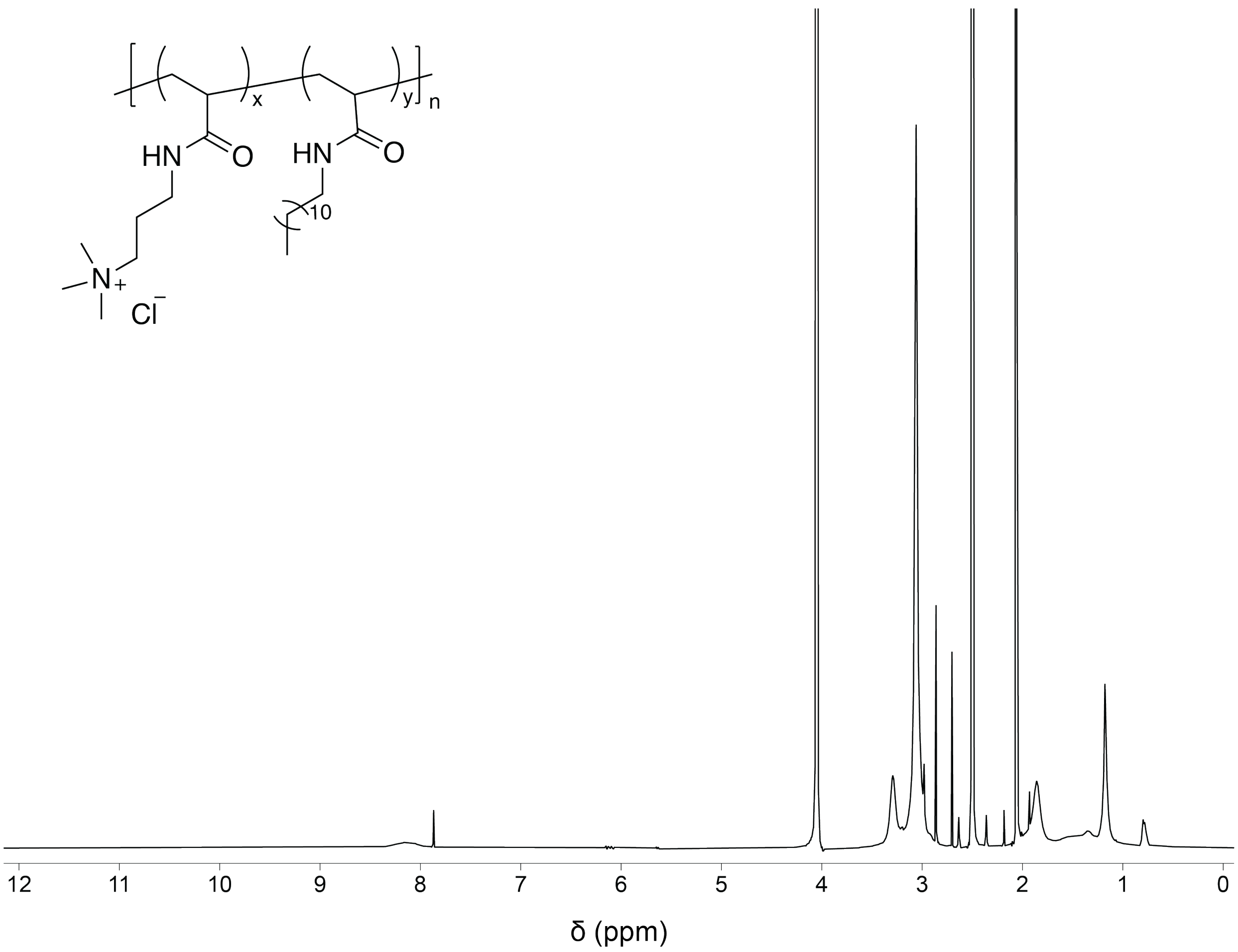

**Figure S4. [1]H NMR Spectrum of Tma$_{75}$Do$_{25}$.** Measured in DMSO-d6/$D_2O$.

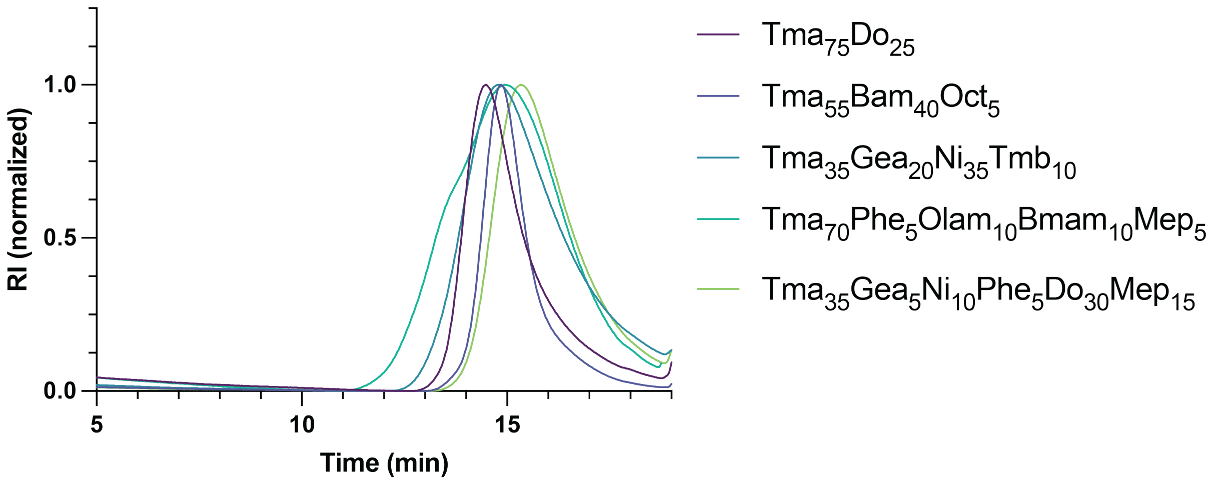

**Figure S5. GPC traces of five most potent candidates.** Each trace was normalized to a minimum value of 0 and a maximum value of 1.

**Table S1. Antibacterial efficacy and safety of novel polyacrylamides.**

| **Polymer** | **MIC: *E. coli*, µg/mL (µM)[a]** | **$HC_{50}$, µg/mL[b]** |
|---|---|---|
| $Gea_{40}Olam_{10}Mo_{50}$ | >512 | 1300 |
| $Gea_{80}Phe_{5}Mo_{15}$ | >512 | <500 |
| $Gea_{5}Phe_{25}Mo_{50}Mep_{20}$ | >512 | 2200 |
| $Tma_{50}Do_{5}Mo_{40}Mep_{5}$ | >512 | >4000 |
| $Tma_{70}Phe_{20}Olam_{5}Mep_{5}$ | 128 | >4000 |
| $Tma_{75}Do_{25}$ | 64 (2.7) | >4000 |
| $Gea_{10}Olam_{30}Mo_{60}$ | >512 | >4000 |
| $Tma_{55}Ni_{25}Phe_{20}$ | >512 | >4000 |
| $Tma_{55}Bam_{40}Oct_{5}$ | 32 (1.7) | <500 |
| $Tma_{17}Phe_{23}Oct_{5}Mo_{55}$ | >512 | >4000 |
| $Tma_{45}Gea_{10}Phe_{20}Olam_{25}$ | 128 | 1500 |
| $Tma_{35}Gea_{20}Ni_{35}Tmb_{10}$ | 64 (3.2) | <500 |
| $Tma_{35}Phe_{35}Olam_{10}Mo_{20}$ | 128 | >4000 |
| $Tma_{38}Gea_{17}Tmb_{8}Mo_{37}$ | 256 | <500 |
| $Tma_{30}Gea_{20}Ni_{20}Olam_{25}Mo_{5}$ | 256 | >4000 |
| $Tma_{90}Do_{10}$ | 256 | >4000 |
| $Tma_{80}Phe_{15}Do_{5}$ | 256 | >4000 |
| $Tma_{85}Gea_{5}Phe_{5}Olam_{5}$ | 128 | >4000 |
| $Tma_{70}Ni_{20}Do_{5}Mep_{5}$ | 256 | >4000 |
| $Tma_{60}Phe_{20}Olam_{5}Mo_{15}$ | 128 | >4000 |
| $Tma_{40}Phe_{5}Do_{5}Mo_{50}$ | >512 | >4000 |
| $Tma_{70}Gea_{10}Oct_{15}Tmb_{5}$ | 192 | <500 |
| $Tma_{65}Do_{5}Bam_{5}Mo_{20}Mep_{5}$ | 256 | >4000 |
| $Tma_{25}Gea_{5}Phe_{15}Bam_{30}Mo_{25}$ | 512 | >4000 |
| $Tma_{70}Phe_{5}Olam_{10}Bmam_{10}Mep_{5}$ | 64 (2.5) | >4000 |
| $Tma_{35}Gea_{5}Ni_{10}Phe_{5}Do_{30}Mep_{15}$ | 64 (5.2) | 3300 |
| $Tma_{30}Do_{15}Bam_{15}Oct_{10}Mo_{30}$ | 128 | >4000 |
| $Tma_{15}Gea_{10}Ni_{5}Do_{25}Tmb_{10}Mep_{35}$ | >512 | <500 |
| $Tma_{25}Ni_{10}Do_{5}Mo_{60}$ | >512 | >4000 |
| $Tma_{20}Gea_{15}Ni_{10}Do_{10}Olam_{10}Mep_{35}$ | >512 | 2500 |

[a]MIC values were determined after overnight incubation with *E. coli*. The µM value was calculated for the five most potent candidates using $M_w$ determined by GPC.

[b]$HC_{50}$ values were determined after 1h incubation with red blood cells.

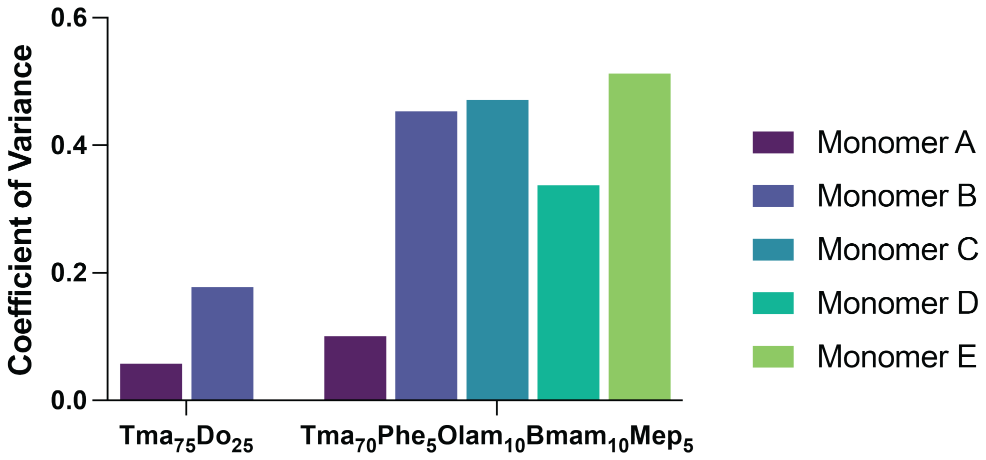

**Figure S6. Compositional dispersity of top-performing candidates.** Calculated using the Monte-Carlo simulation, Compositional Drift. This result demonstrates higher compositional variance for the 5-monomer system compared to the 2-monomer system.

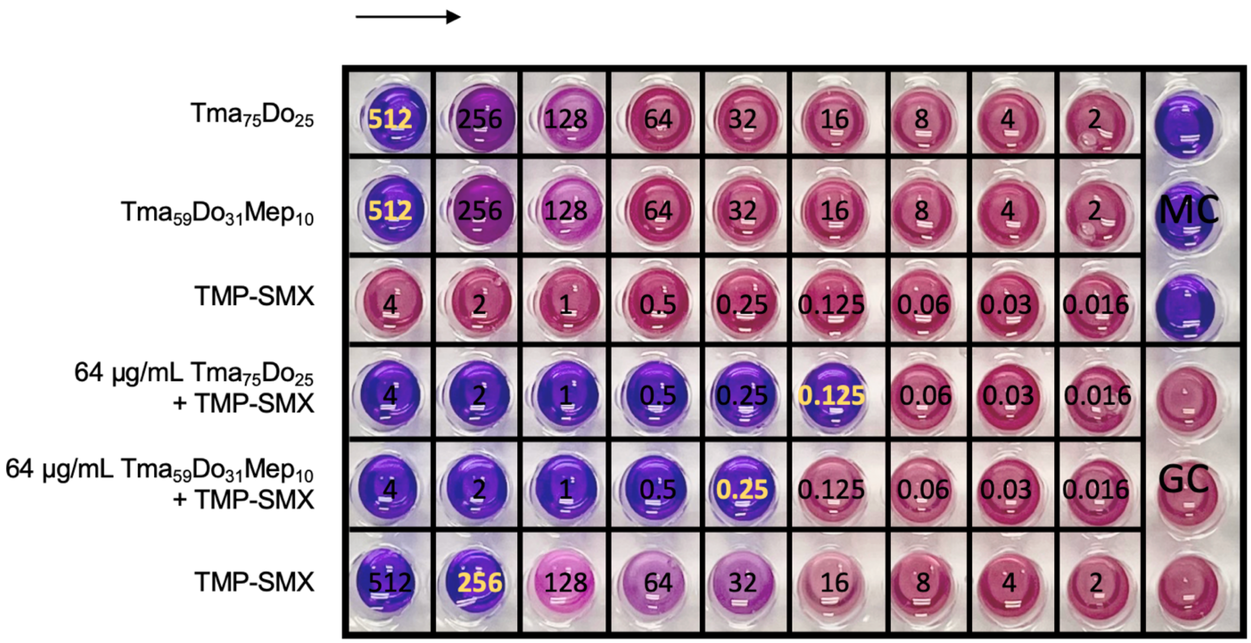

**Figure S7. MBEC assay of lead candidates and TMP-SMX**, alone and in combination. Developed with resazurin dye. Yellow values highlight the MBEC for each compound in µg/mL.
